# Neural Sharpening of Object Categories Through Parafoveal Priming

**DOI:** 10.1101/2025.05.16.654463

**Authors:** Syanah C. Wynn, Yulia Bezsudnova, Ole Jensen

## Abstract

During natural vision, humans make saccades approximately every 250 - 300 ms to fixate on important objects in a scene. Given this rapid succession of eye movements, it has been proposed that visual processing begins in the parafovea, prior to fixation. Such parafoveal previewing may serve as a form of priming, facilitating subsequent processing at fixation. In this study, we investigated whether parafoveal priming leads to both a reduced neural response - akin to repetition suppression - and a more selective neural representation, consistent with neuronal sharpening. Participants performed a visual exploration task involving natural object images that were either parafoveally previewed or not, while magnetoencephalographic (MEG) data were recorded. Event-related fields revealed a reduced neural response when an image had been parafoveally presented 250 ms earlier. Moreover, multivariate pattern analysis showed enhanced decoding of object category following parafoveal priming, suggesting increased neural specificity. These findings support the idea that parafoveal previewing contributes to neural sharpening, potentially aiding evidence accumulation by forming sparse and precise representations of objects during visual scene exploration.

## 1. Introduction

When exploring visual scenes, we make rapid eye movements (saccades) to fixate on important or relevant objects. Saccades are initiated every 250 – 300 ms bringing objects from the parafovea into the fovea, where they can be processed with higher visual acuity. As about 100 ms are needed to initiate and execute each saccade, this leaves only 150 – 200 ms to process foveal information before the eyes move to the next object ^1,2^. It has been proposed that visual processing can be given a head-start by parafoveally processing objects before eye movements are made to them ^1-4^.

The preview benefit associated with parafoveal processing can be considered a form of priming, as stimuli are viewed repeatedly, first parafoveally and then foveally. While this stimulus repetition may have behavioural benefits, repetition is also associated with a reduction in the neural response, an effect referred to as repetition suppression ^5,6^. When measured using electroencephalography (EEG), repetition suppression occurs between 150 and 200 ms after stimulus onset over posterior regions ^7^. The question of how a reduction in the neural response relates to the behavioural preview benefit may be answered by the concept of “sharpening” ^8-11^. The concept of sharpening refers to engaging only the population of neurons coding for the key features of an object thereby narrowing the respective tuning curves. For instance, when a *sunflower* is shown, neurons representing flowers in general might initially fire. However, with sharpening, neurons only coding for a *sunflower* might activate and the neuronal response thereby becomes more selective. The sharpening might occur with prolonged or repeated exposures to a given object - or when the object can be predicted from recent history ^12^. Sharpening may also account for the decrease in the population’s neuronal response with repetitions. It has been proposed that synchronization in the gamma-band frequency facilitates the inhibitory mechanisms associated with sharpening ^13,14^. This follows from the finding that repetition of images elicits both a reduction in neuronal firing rates and an initial dip followed by a subsequent increase in 40-80 Hz power in the macaque ^13,14^ and human visual cortex ^15^.

The aim of the current study is to investigate whether parafoveal priming is accompanied by repetition suppression as well as sharpening. To this end, we presented images of natural objects foveally that either were primed by the same preceding images in the parafovea or not. To investigate the neuronal mechanisms associated with parafoveal priming, we utilized magnetoencephalography (MEG) to record brain activity. We used the MEG data to compute event-related fields (ERFs), gamma power, as well as to decode the object category using a support vector machine (SVM) classifier. MEG enabled us to investigate the time course of the parafoveal priming at a millisecond time scale investigating both evoked responses as well as using multivariate pattern analysis (MVPA). Our specific hypotheses were that the parafoveal priming would result in: (1) repetition suppression, demonstrated by a reduction in the ERF between 150 and 200 ms after foveal stimulus onset; (2) sharpening, demonstrated by an increase in object category classification performance and possibly a decrease in gamma power in the same time window as the repetition suppression.

## 2. Methods

### 2.1. Participants

Thirty-five healthy adult volunteers (23 females, 12 males) with a mean age of 21.83 (SD = 3.30) participated in this study. All had normal or corrected-to-normal vision, were fluent English speakers, and free from self-reported neurological or psychiatric conditions. Main exclusion criteria were metal implants, diabetes, asthma, claustrophobia, attention deficit (hyperactivity disorder (ADHD), autism spectrum disorder (ASD), epilepsy, other neurological/psychiatric conditions, and pregnancy. The study was approved by the institutional review board of the University of Birmingham and carried out in accordance with the standards set by the Declaration of Helsinki. All participants received written and oral information prior to participation but remained naive regarding the aim of the study. Each participant provided written informed consent at the beginning of the first session.

### 2.2. Stimuli

Stimuli were obtained from the freely available THINGS database, which consists of multiple naturalistic images of almost two thousand object ^16^. From this database, objects were selected according to the five categories: animals, clothing, food, plants, and vehicles. Within each of these object categories, one hundred objects were picked and for each object, fifteen images were selected. This led to a total of 7500 unique stimuli (5 object categories × 100 objects × 15 images). When the THINGS database was not sufficient to provide enough images, additional images were obtained through Pl@ntNet, Pixabay, and Unsplash databases. All images were cropped to 480 × 480 pixels in MATLAB (v2021a, MathWorks Inc., Natrick MA) and presented by a PROPixx DLP LED projector (VPixx Technologies, Canada). Stimulus presentation and recording of responses were attained using the MATLAB Psychophysics Toolbox Version 3 (PTB-3) ^17-19^.

### 2.3. Experimental paradigm

The experimental paradigm was designed to mimic naturalistic viewing in a controlled fashion (see Figure 1). During naturalistic viewing, objects may initially appear in the parafovea followed by a foveal fixation. In the current experiment, natural viewing was emulated by having participants fixate the gaze in the middle of the screen while objects were presented in the parafovea and the fovea. In this way, we controlled the visual information presented to the participants while avoiding saccade producing artifacts in the MEG signal. The experimental task consisted of 12 blocks, composed of a “viewing” and a “memory” phase. Each block consisted of 120 trials in the viewing phase and 60 trials in the memory phase, totalling 1440 viewing and 720 memory trials in the full experiment.

**Figure 1.**
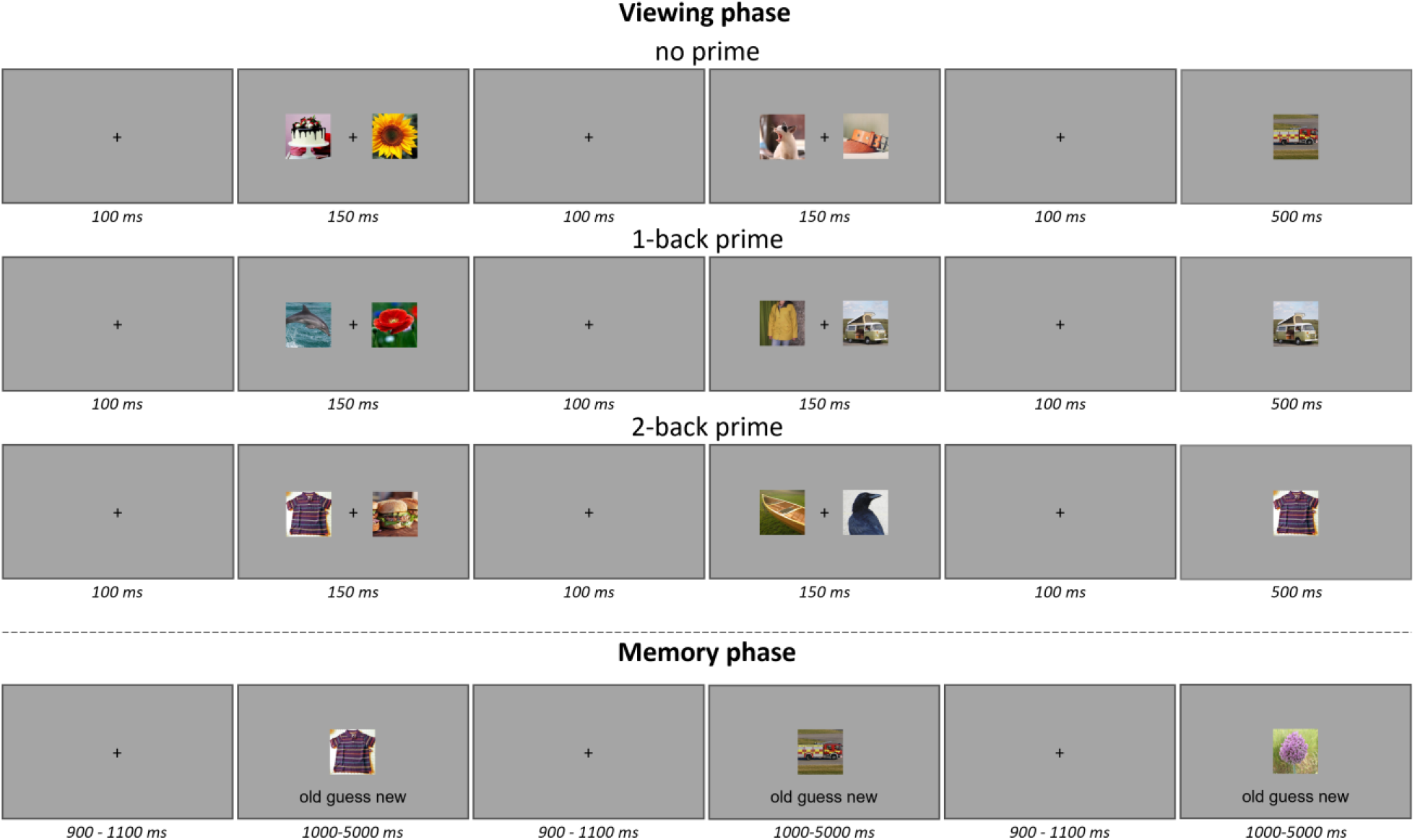
Experimental paradigm. In the viewing phase (top) there were three types of trials. All trials consisted of two consecutive displays of two simultaneously presented parafoveal objects followed by a single object presented foveal. In the “no prime” condition, the foveal object was not primed prior to presentation. In the “1-back prime” condition, the foveal object was primed parafoveal one display back. In the “2-back prime” condition, the foveal object was primed parafoveal two displays back. In the memory phase (bottom) consecutive objects were presented centrally. Half of these object were presented foveal in the viewing phase (“old”) and half of these object were not presented before (“new”). Participants were instructed to indicate whether the object was “old” or “new”. The memory phase was self-paced, with the constraint that each object was presented for a minimum of 1000 and a maximum of 5000 ms.

The viewing phases started with a 100 ms central fixation cross, followed by the presentation of two objects in the left and right parafovea for 150 ms. After a 100 ms central fixation cross followed the presentation of another two parafoveal objects for 150 ms, a 100 ms central fixation cross, and finally the presentation of a foveal object for 500 ms (see Figure 1). All images were scaled to have the dimensions of 15% of the height of the screen. The experiment was projected on a 71 × 41 cm screen, resulting in the size of the images being 6.15 × 6.15 cm. The viewing distance between the screen and the participant was 145 cm. Foveal objects were presented in the centre of the screen and parafoveal objects were presented on the left and right sides of the centre of the screen. The foveal image had a visual angle of 2.4°, centred at 0°. The parafoveal images spanned a visual angle from 2.3° to 7.2°, with the centre of the image being at a visual angle of 4.8°.

There were three types of trials (see Figure 1): trials where the foveal object was not presented before (“no prime”), trials where the foveal object was also previously presented parafoveally just before foveal stimulus onset (“1-back prime”), trials where the foveal object was also presented parafoveally at the start of the trial (“2-back prime”). Therefore, in the latter two conditions, the foveal object was primed parafoveally either one display back or two display back by presenting the identical images parafoveally.

Regarding the foveal images, every object was shown once per priming condition, through random selection of a unique image representing the object. Therefore, every trial consisted of unique images which were not repeated over trials. Each of the twelve blocks contained an equal number of images per category and priming condition while ensuring no more than two consecutive identical priming conditions and no more than four consecutive identical object categories. Due to these balancing requirements, 96 out of the 100 available objects were randomly chosen to be shown foveally per participant.

In the “1-back prime” and “2-back prime” conditions, the foveal image was also presented parafoveally, with an equal probability of being shown on either the left or right side of the screen. These priming images were counterbalanced to ensure each object category had an equal chance of appearing in each parafoveal position (i.e., 1-back left, 1-back right, 2-back left, 2-back right).

For the “no prime” condition, the parafoveal images consisted of one random image from each of the four object categories that remained after excluding the object category of the foveal image. In the “1-back prime” and “2-back prime” conditions, the parafoveal images included one random image from three out of the four object categories that remained after foveal object category exclusion. These images were randomly assigned over the available parafoveal positions.

The memory phases started with a central fixation cross that had a jittered duration (range: 900-1100 ms). After, an image was presented centrally with the response options (“Old”, “Guess”, “New”) at the bottom of the screen. The memory task was self-paced, with the constraint that each image was presented for a minimum of 1000 and a maximum of 5000 ms. Participants were instructed to indicate if an image was “Old” when they had seen it earlier, “New” if they had not seen it before, and “Guess” if they were not sure whether they had seen it or not. They were instructed to base their decisions on all the images seen in the viewing phase of the current block. Participants used their right index, middle, and ring finger to indicate their responses using a button box. The “Old” and “New” options were randomly mapped to the index finger for half the participants and the ring finger for the other participants. The “Guess” option was always mapped to the middle finger. Participants were informed about their memory performance at the end of each block.

For each block, the 30 “Old” images were randomly picked from the 120 foveal images of the current block. The 30 “New” images were chosen from the pool of images not presented in any of the blocks. To ensure a representative sample, they were balanced for object category and priming condition (“Old” images), or object category only (“New” images). The order of presentation of these “Old” and “New” images was randomized.

Participants were familiarized with the task through a practice block, which contained 15 viewing trials and 10 memory trials. To aid in the understanding of the task, immediate feedback was given after the memory trials by presenting “correct!” or “incorrect!” to the participants. Participants could repeat the practice block until they felt confident in their understanding of the task. None of the images in the practice block were used in the main experimental blocks.

### 2.4. Data acquisition

#### 2.4.1. Eye movement data

An EyeLink 1000 Plus eye tracker (SR Research, Canada) was used to continuously record eye movement data throughout the experiment. The eye tracker was positioned on a table 90 cm in front of the participant. At the beginning of the experiment, a 9-point calibration and validation procedure were conducted. Additionally, a drift check was performed after each trial to correct for any linear drift. Eye movement data, including the x and y positions and pupil size of the left eye, were captured at a sampling rate of 1000 Hz.

#### 2.4.2. MEG data

MEG data were recorded using a 306-sensor TRIUX MEGIN system, consisting of 204 orthogonal gradiometers and 102 magnetometers, with an online band-pass filter of 0.1 to 330 Hz and a sampling rate of 1000 Hz. Before data collection, four head-position indicator (HPI) coils were attached to the head of the participant: two on the forehead, with a minimum separation of 3 cm, and two on the left and right mastoid bones behind the ears. Using the Polhemus Fastrack electromagnetic digitizer system (Polhemus, USA), we first recorded the positions of three anatomical fiducial points: the nasion, and the left and right preauricular points. We then digitized the positions of the four HPI coils and at least 300 points on the scalp to capture the head shape. Participants were then seated in the MEG gantry (60 degrees upright position) for the actual experiment.

### 2.5. MEG preprocessing

Data were analysed using the open-source toolbox MNE Python v1.4.2 ^20^ according to the standards of the FLUX pipeline ^21^. Only MEG data pertaining to the viewing phase are presented here.

The eye-tracking data were used to add markers to the MEG data for the occurrence of blinks, saccades, and gaze fixation. The time points and type of occurrence were automatically detected through the Eyelink software. In addition, eye-tracking data were used to emulate an electrooculography (EOG) channel.

The initial step of the preprocessing differs for the univariate and multivariate analyses. For the univariate analysis, a Maxwell filter for Signal Space Separation (SSS) and noise reduction was applied and the continuous HPI (cHPI) recording was used to correct for head motion. For the multivariate analyses we did not perform the SSS first as this might be suboptimal for multivariate analysis ^22^. In the preprocessing we opted to interpolate bad sensors. The following preprocessing steps were identical for univariate and multivariate analyses.

Data segments which included muscle artefacts, where the eyes were closed for more than one second, and blinks in the first 100 ms of stimulus onset (parafoveal and foveal) were automatically marked as “bad” in the MEG data. These “bad” data segments were ignored in future preprocessing steps and later rejected. Independent component analysis (ICA) was used to suppress eye movement and heartbeat-related artefacts. The ICA components were estimated after the data were bandpass filtered at 1 – 40 Hz and down-sampled to 250 Hz. Eye-movement and heartbeat-related components were automatically detected through the correlation of the independent components with the emulated EOG and Electrocardiography (ECG) channels using a correlation threshold of 0.2. The automatically selected eye-movement and heartbeat-related components were then manually checked for each participant and corrected when necessary. The final selection of components was used to attenuate the EOG and ECG artefacts in the original (unfiltered; non-down sampled) data. The resulting data were then bandpass filtered (0.1 – 150 Hz) and detrended before it was epoched from -1500 to 1500 ms, relative to the foveal stimulus onset. At this stage, the trials with annotated artefacts were excluded. In addition, trials with excessive peak-to-peak values were excluded according to the FLUX guidelines. Participants with less than 50% of trials left were removed from further analyses. This led to 31 participants to be considered for the univariate analyses and 32 participants to be considered for the multivariate analysis.

### 2.6. Behavioural analysis

Memory accuracy per priming condition (“no prime”, “1-back prime”, “2-back prime”) was quantified by d-prime, according to the following formula:

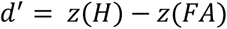

Where *z(H)* is the z-transform of the hit rate and *z(FA)* the z-transform of the false alarm rate. Only participants that were included in the univariate and/or multivariate analysis were included in the behavioural analysis, leading to a total of 33 participants in this analysis. Differences in priming conditions (within-subject factor) were statistically tested with a repeated measure analysis of variance (RM ANOVA) and follow-up t-tests comparing “no prime” trials to “1-back prime” and “2-back prime” conditions.

### 2.7. MEG analysis

#### 2.7.1. Sensor selection for univariate analyses

Sensor selection for the ERF and spectral (but not the multivariate) analyses was based on visual evoked fields (VEFs) in the grand average in the 0 – 500 ms time window. A cluster of the sensors showing the greatest VEF was chosen separately for the magnetometers and gradiometers (see Figure 2). The VEFs for the magnetometers and the gradiometers were considered separately when identifying the sensors of interest.

**Figure 2.**
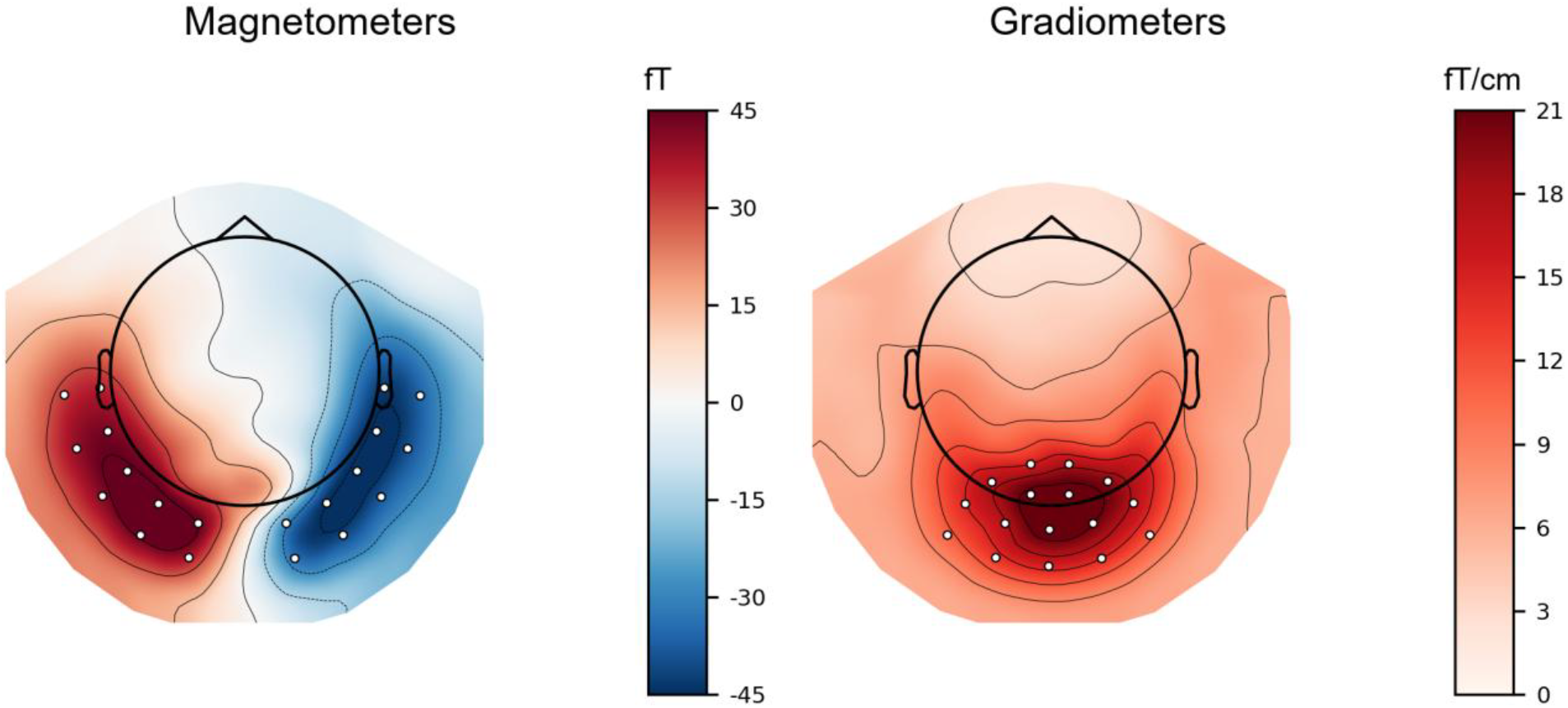
Grand average of the visual evoked responses (0 - 500 ms). The magnetometer (left) and gradiometer (right) sensors selected for the ERF and spectral analyses indicated in white.

#### 2.7.2. Event-related fields (ERFs)

Prior to evaluating the task-dependent ERFs, the data were low-pass filtered at 30 Hz, and then a baseline subtraction was applied using the 100 ms prior to the foveal stimuli onset ^21^. ERFs were then computed by averaging the signal over trials for each sensor. The planar gradiometers were combined by calculating the root mean square of orthogonal gradiometer pairs with the same location.

Cluster-based permutation (CBP) was used to test for statistically significant effects in the ERFs. CPB was performed separately on gradiometers, left magnetometers, and right magnetometers. CBP was applied in the time window of the foveal image presentation (0 – 500 ms) and the predefined MEG sensor selection. The CBP compared “no prime” to the “1-back prime” and “2-back prime” conditions. An alpha value of .005 determined the cluster-forming threshold for the two-sided t-tests. This threshold was used because it reduces the influence of large diffuse clusters with weaker effects. Given that we expected the effect to be focal in time, we opted to use a more conservative threshold for the forming of the clusters. A p-value of .05 was subsequently used to determine if these clusters reflected a significant difference, based on the permutation test.

#### 2.7.3. Multivariate Pattern Analysis (MVPA)

The data were first epoched (−850 – 650 ms; t = 0 defines the onset of the foveal image) and all MEG sensors were included in the analysis. A standard SVM classifier with a linear kernel and balanced weights was used to classify the object categories. Prior to applying the classification algorithm, the data from each sensor was z-scored. We utilized temporal decoding, which fits a multivariate predictive model on each time point and evaluates its performance at the same time point on new data. The classifier was trained on 80% of the data and tested on the remaining 20% (5-fold cross-validation). The outcome measure was the area under the curve (AUC). The mean AUC across the five cross-validation splits was used as the classification score. Classification is done using the one versus others approach for each priming condition (“no prime”, “1-back prime”, “2-back prime”). Specifically, every object category within a specific priming condition is compared against the other four categories across priming conditions. This leads to an unbiased and comparable classification across the priming conditions. As there were five categories in total, this provided five classification scores. These five classification scores were averaged per priming condition, leading to a single combined classification score per priming condition.

CPB was used to compare the “no prime” to the “1-back prime” and “2-back prime” conditions in the 0 – 500 ms interval on all channels. The CBP compared “no prime” to the “1-back prime” and “2-back prime” conditions. An alpha value of .005 determined the cluster-forming threshold for the two-sided t-tests. A p-value of .05 was subsequently used to determine if these clusters reflected a significant difference, based on the permutation test.

#### 2.7.4. Time-frequency representation (TFR)

Spectral power of the data was extracted using Fourier analysis with a sliding time window of 250 ms and a frequency smoothing of ± 8 Hz, through the application of multitapers (3 Discrete Prolate Spheroidal Sequences (DPSS) tapers). Frequencies were assessed from 30 to 100 Hz in 2 Hz steps ^21^.

CBP was used to test statistically significant effects in the time-frequency representations of power. Analyses were performed separately on gradiometers, left magnetometers, and right magnetometers. Data were averaged across the MEG sensors that showed the largest VEFs (see above). CBP focused on the time window of the foveal image presentation (0 – 500 ms) and on the frequency range of 40 to 80 Hz. The CBP compared “no prime” to the “1-back prime” and “2-back prime” conditions. An alpha value of .005 determined the cluster-forming threshold for the two-sided t-tests. A p-value of .05 was subsequently used to determine if these clusters reflected a significant difference, based on the permutation test.

## 3. Results

Participants viewed two sets of images presented in the left and right hemifield after which an image was presented in the fovea (Figure 1). This resulted in three parafoveal priming conditions (“no prime”, “1-back prime”, “2-back prime”). The memory for these images was subsequently tested. The brain activity was recorded throughout using MEG, and behavioural and neuronal effects are reported below.

### 3.1. The 1-back priming effect improves memory

The analysis showed a significant difference in memory performance across priming conditions (F(2,64) = 4.25, p = 0.02; see Figure 3). A follow-up t-test showed that there was improved memory performance for the “1-back prime” (M = .66) compared to the “no prime” (M = .58) condition (t(32) = 2.28, p = 0.03). There was no difference between the “2-back prime” (M = .57) and “no prime” conditions (t(32) = -0.10, p = 0.92). These findings suggest that the parafoveal presentation of the object just before the foveal presentation enhanced memory performance. This effect disappeared when an intervening object was presented between the parafoveal prime and the foveal target.

**Figure 3.**
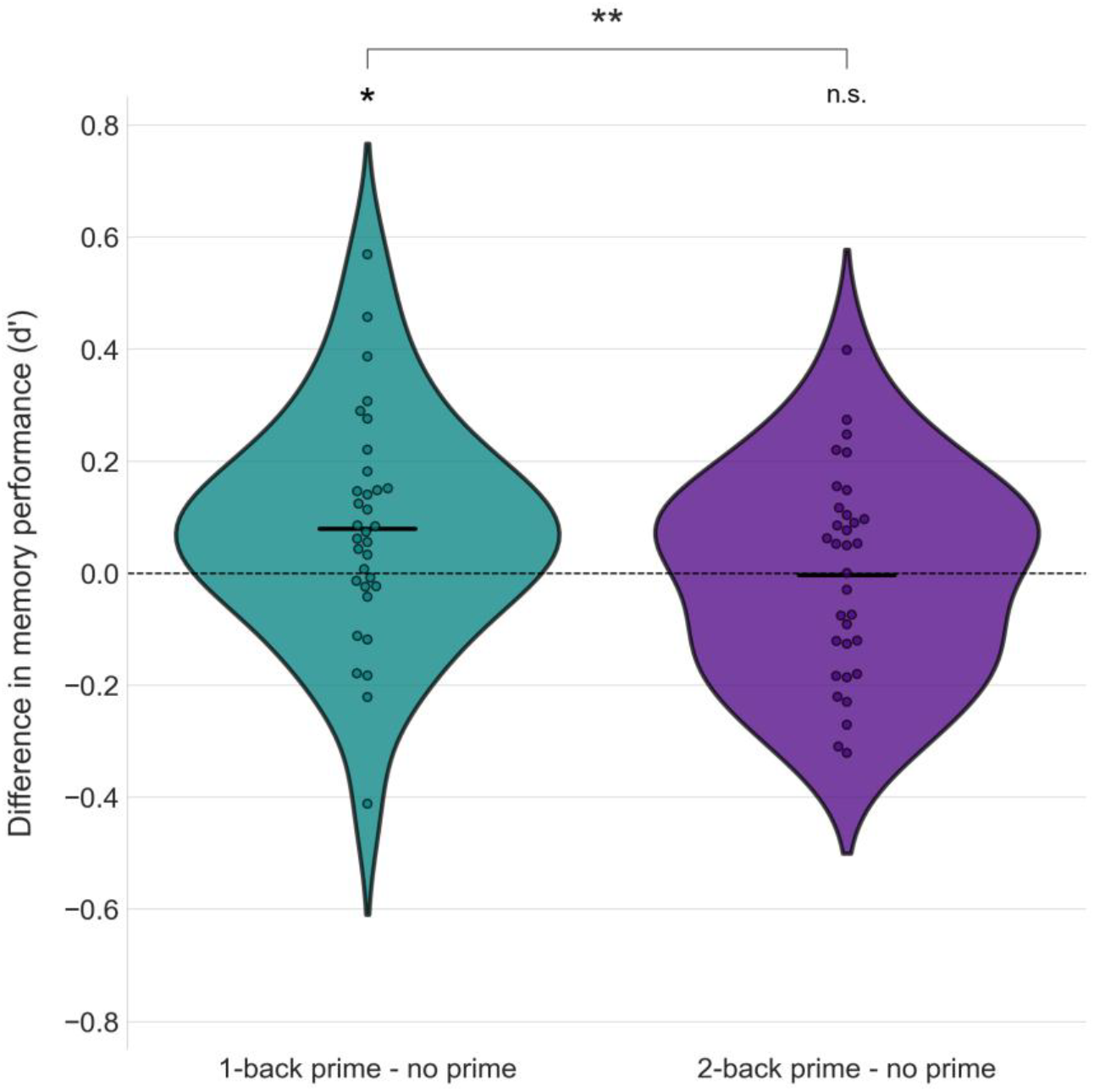
Difference in memory performance. Quantified as d-prime (d’) difference between the “1-back prime” and “no prime” conditions (left; t(32) = 2.28, p = 0.03) and the “2-back prime” and “no prime” conditions (right; t(32) = -0.10, p = 0.92). The difference between the ‘1-back prime’ and ‘no prime’ conditions was significantly larger than the difference between the ‘2-back prime’ and ‘no prime’ conditions (t(32) = 2.78, p = 0.009). Asterisks indicate statistical significance: *p<0.05, **p<0.01.

### 3.2. Repetition suppression observed in the ERFs

When analysing the ERF in the sensors with a strong visual evoked response (see Figure 2), we observed a reduced response for the “1-back prime” compared to the “no prime” condition in the selected sensors (all *p*s < .03, see Figure 4). This difference was most pronounced in the 150 – 210 ms interval. In addition, there was a significant difference between the “no prime” and “2-back prime” conditions in the left magnetometers (*p* = .010). This difference was most pronounced in the 190 – 210 ms interval. These findings provide evidence for repetition suppression, i.e. a diminished response at ∼175 ms, when an object was presented parafoveally just before the foveal presentation. The repetition suppression was reduced for the “2-back prime” condition. Furthermore, when considering the later part of the ERF, we observed an increased response for the “1-back prime” compared to the “no prime” condition in the right magnetometer (*p* = .002), which was most pronounced in the 305 – 360 ms interval. This may reflect repetition enhancement at ∼325 ms, when an object was presented parafoveally just before the foveal presentation. There was no significant repetition enhancement for the “2-back prime” condition. In sum, the 1-back prime results in a robust repetition suppression as reflected in the ERFs at ∼175 ms.

**Figure 4.**
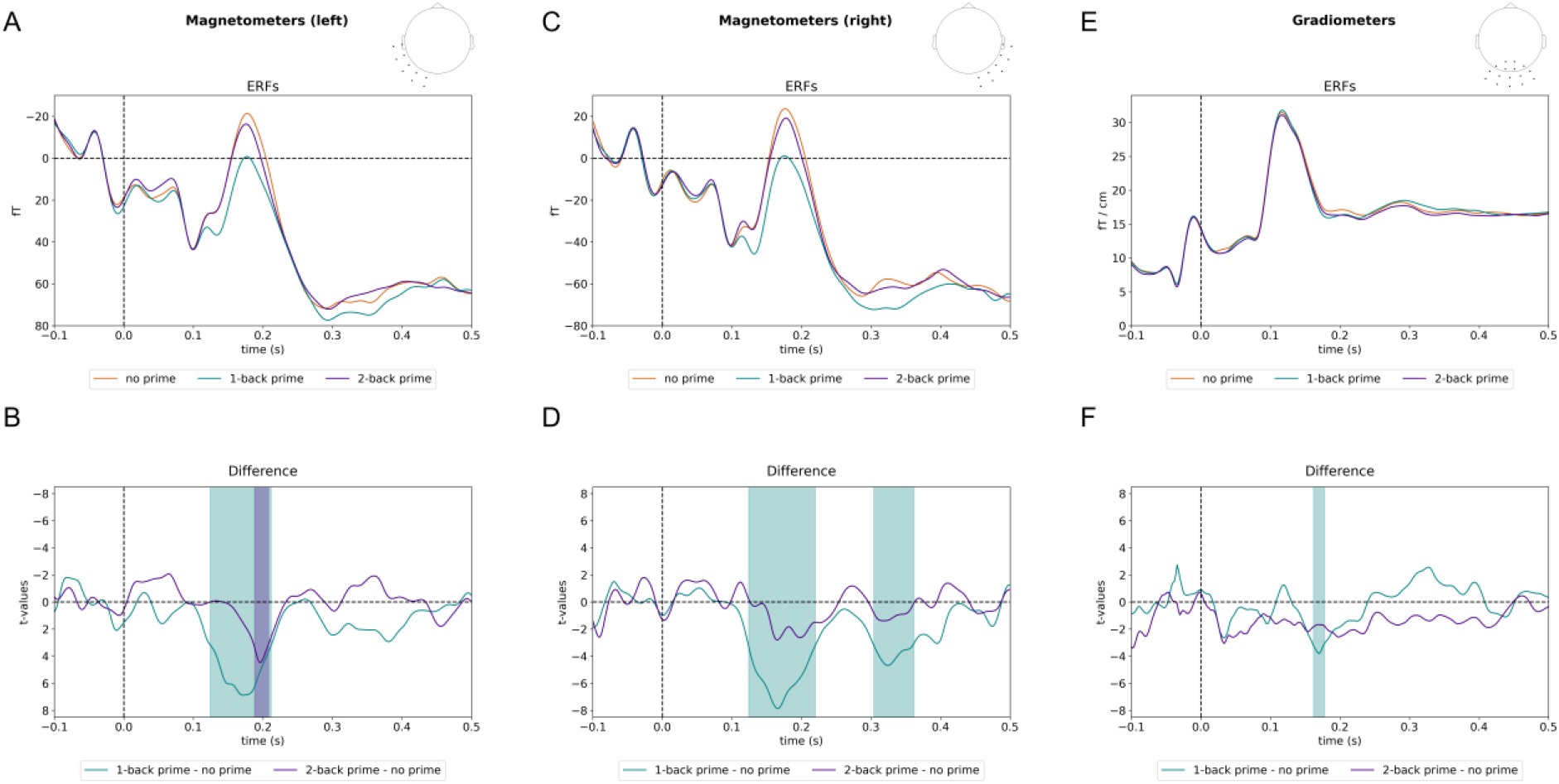
Event-related fields (ERFs) for the three priming conditions (no prime, 1-back prime, 2-back prime) over the three sensor clusters. (A) ERFs for the left magnetometers. Note that due to difference in polarity between left and right sensors, the axis for the left ERF a plotted with negative up. (B) The t-values of the difference between 1-back prime and no prime, and 2-back prime and no prime conditions, with the significant clusters shown in the respective coloured area (p = .001 and p = .01, respectively). (C) ERFs for the right magnetometers. (D) The t-values of the difference between 1-back prime and no prime, and 2-back prime and no prime conditions, with the significant clusters shown in the respective coloured area (early: p = .001 and late: p = .002). (E) ERFs for the gradiometer sensors. (F) The t-values of the difference between 1-back prime and no prime, and 2-back prime and no prime conditions, with the significant clusters shown in the respective coloured area (early: p = .005 and late: p = .03).

### 3.3. Improved decoding with parafoveal priming

The classification analysis shows that we could robustly decode the object category of the foveally presented image with increasing accuracy from 100 ms onwards (see Figure 5A). The classification accuracies were significantly different between the “1-back prime” and “no prime” conditions (p = .016), and between the “2-back prime” and “no prime” conditions (p = .009). These differences were most pronounced in the 355 – 380 ms and 415 – 440 ms time intervals, respectively (see Figure 5B).

**Figure 5.**
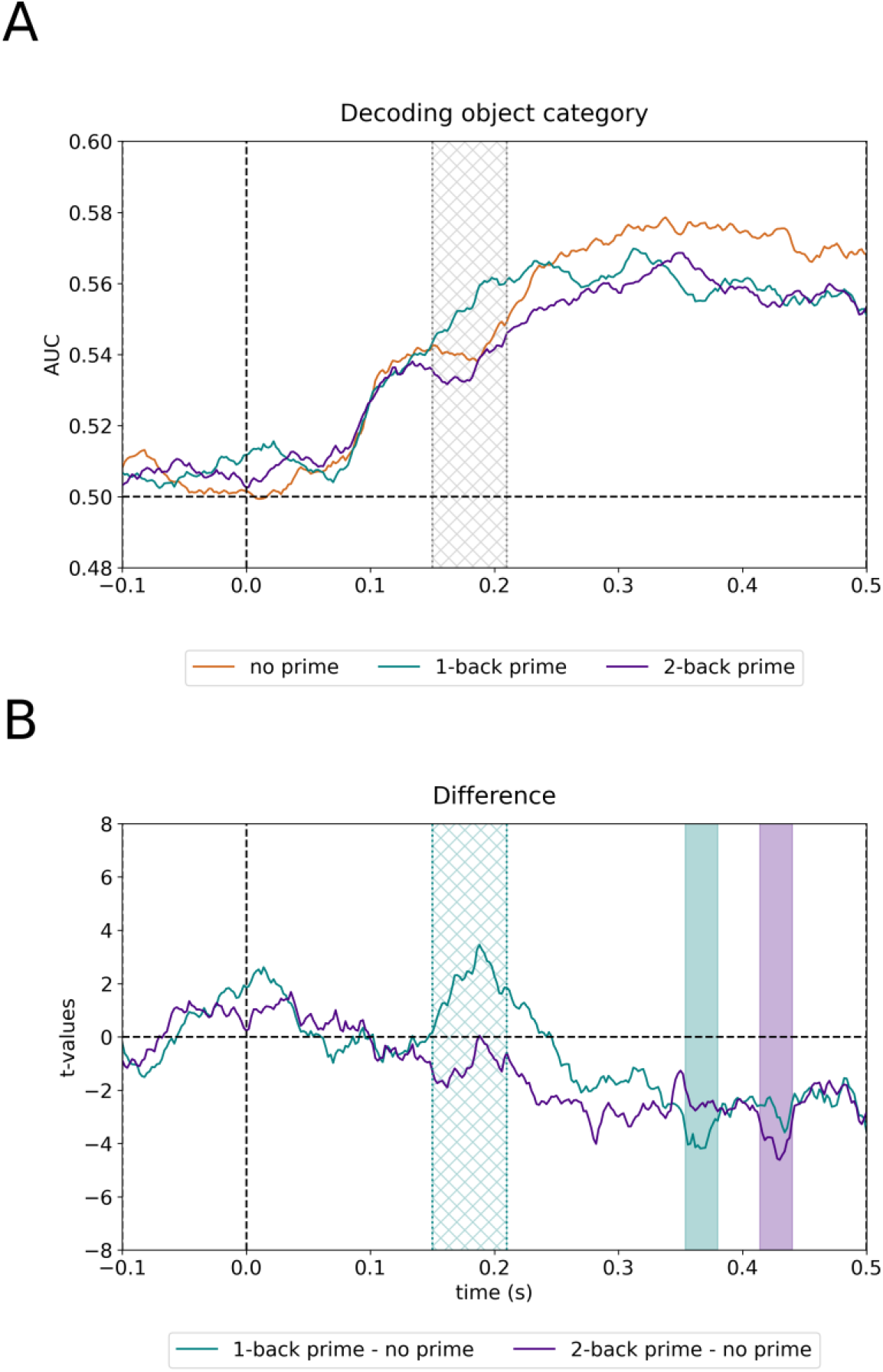
Decoding of the category of the foveal image. (A) Classification scores per priming condition (no prime, 1-back prime, 2-back prime) with 0.50 representing chance level. (B) The t-values of the difference between 1-back prime and no prime, and 2-back prime and no prime conditions, with the significant clusters shown in the respective filled-in coloured area (p = .016 and p = .009, respectively). The cross-hatched area represents the time window of repetition suppression (150 – 210 ms) in which our post-hoc tests showed a significant increase in classification accuracy for the “1-back prime” as compared to the “no prime” conditions (t(31) = 2.55, p = 0.02).

Given the specific hypothesis regarding sharpening in the interval where we also observed repetition suppression (150 – 210 ms; see ERF results), we conducted a post-hoc repeated measures ANOVA with condition (no prime, 1-back prime, 2-back prime) as the within-subject factor. This revealed a significant difference between conditions (F(2,62) = 9.31, p < 0.001). Follow-up t-tests showed that there was an increased classification accuracy for the “1-back prime” (M = .55) compared to the “no prime” (M = .54) condition (t(31) = 2.55, p = 0.02). There was no significant difference between the “2-back prime” (M = .54) and “no prime” conditions (t(32) = -1.42, p = 0.17). This shows that when considering the repetition suppression time window, there was evidence for improved decoding when priming parafoveally, but only for the 1-back prime condition.

### 3.1. No robust support for sharpening reflected by modulation in gamma oscillations

When analysing the spectral data in the posterior sensors with a strong visual response, we observed no robust difference in gamma power during the “1-back prime” nor the “2-back prime” condition, as compared to the “no prime” condition (*p*s > .204; see Figure 6). Nevertheless, when visually inspecting the time-frequency representations of power differences, there was a tendency for the priming conditions to result in stronger 50-70 Hz gamma power; this effect was however not statistically significant. Our finding does therefore not show robust evidence for modulation in gamma activity when an object was parafoveally primed.

**Figure 6.**
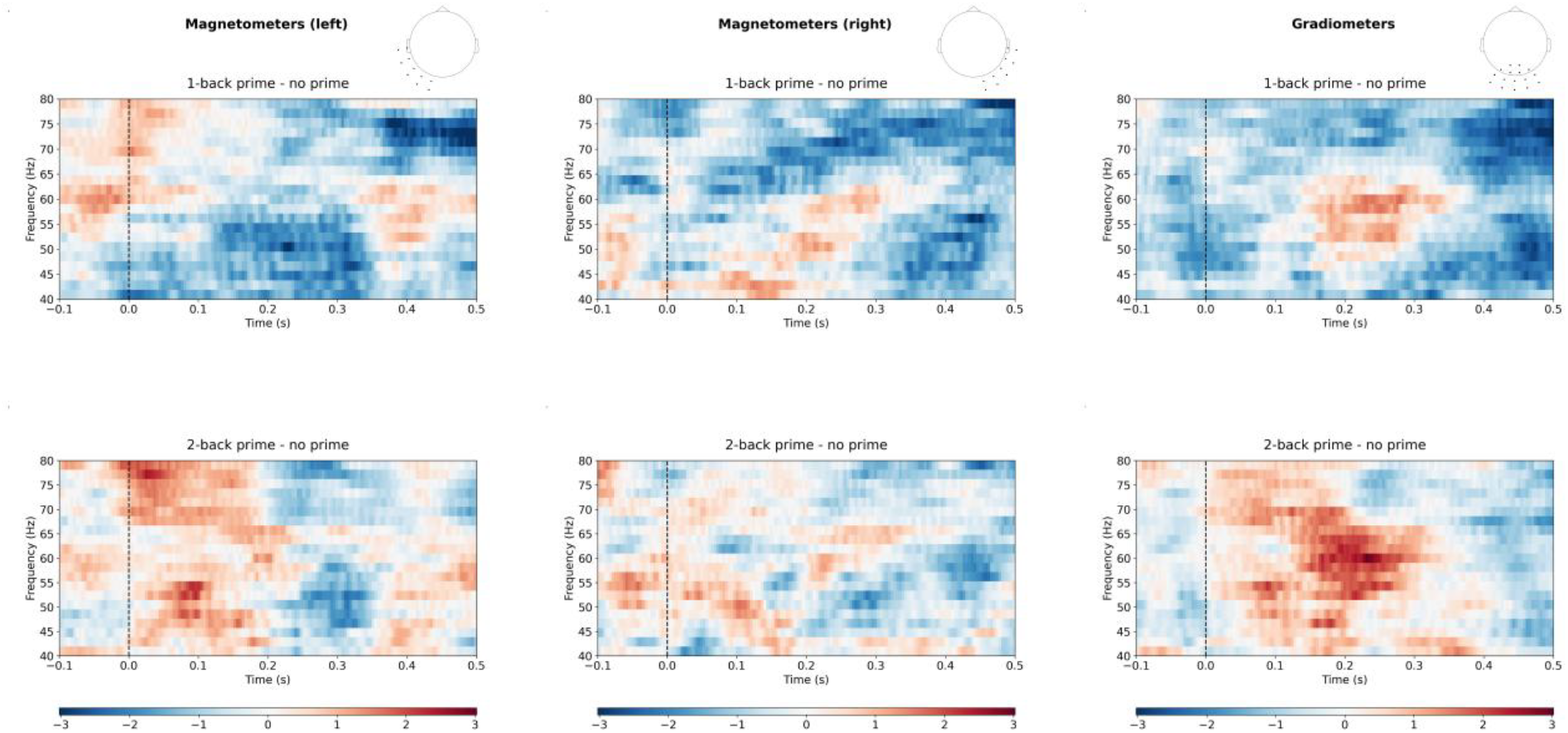
Time-frequency representations (TFRs) of gamma power differences. Gamma power differences are shown for “no prime” versus “1-back prime” and “no prime” versus “2-back prime” conditions. (A) shows the t-values for the gamma differences between “no prime” and “1-back prime” for the left magnetometers, while (B) shows the same comparison for “no prime” and “2-back prime.” (C) and (D) display the t-values for these conditions for the right magnetometers. (E) and (F) present the t-values for the “no prime” vs. “1-back prime” and “no prime” vs. “2-back prime” comparisons for the gradiometers, respectively. The t-values do not research significance when the multiple comparisons over time-and frequency are considered.

## 4. Discussion

The current study explored whether parafoveal previewing is associated with sharpening by investigating whether parafoveal priming is accompanied by both repetition suppression and a more specific neural code. To this end, our participants completed a task where foveal images were primed parafoveally while we recorded their brain activity with MEG. Our results suggest that parafoveal previewing is indeed associated with sharpening. This is reflected by the parafoveal priming-induced repetition suppression in the ERFs, time-locked to foveal image onset, within the 150 – 210 ms time window. Importantly, we found evidence for improved decoding of object categories after parafoveal priming, indicated by increased classification accuracy. Lastly, we found no evidence for the hypothesis that synchronization in the gamma-band frequency facilitates the inhibitory mechanisms in sharpening.

### 4.1. Parafoveal priming and sharpening as an accumulation of evidence process

In natural settings our eyes are moving from object to object, a process typically resulting in both parafoveal and then foveal processing of a given object. Our findings suggest that objects in the parafovea can be identified at the category level as reflected by the multivariate analysis of the MEG data. This parafoveal processing speeds-up of the categorisation of the object when fixated. We propose that parafoveal followed by foveal processing reflects the accumulation-of-evidence resulting in a more precise distributed neuronal code involving less neurons. This mechanism supports natural viewing as the parafoveal previewing serves to give visual processing a head-start and procure a more accurate and sparse neuronal representations. In future work it would be of interest to explore computational neuronal models that can account for this mechanism. Key components of such a model could involve recurrent connections between neurons representing a given object with short-term Hebbian synaptic strengthening in combination with lateral inhibition reducing the activation of neurons receiving input below a certain threshold.

### 4.2. The behavioural benefit of parafoveal priming is associated with repetition suppression

It is well-established that priming can have an influence on behaviour, including the facilitation of object recognition ^23-26^. In line with this, our behavioural results show that participants were significantly better at remembering the images when parafoveal priming directly preceded the image. This suggests that parafoveal priming facilitated the encoding of the object that was subsequently presented. We furthermore showed that parafoveal priming was associated with repetition suppression. Specifically, we found a reduction in the ERF when the image was parafoveally primed right before the foveal image, as compared to when no priming occurred, in the 150 – 210 ms time-window. These results closely align with other event-related repetition suppression literature ^7,27-29^. While previous studies presented visual objects at a single location, we extend these findings by demonstrating similar repetition suppression effects when priming occurs parafoveally. This means that the priming mechanism must occurs in areas where the parafoveal and foveal representations overlap in terms of receptive fields, i.e. at the earliest in V4. The results described here suggest a partial processing of parafoveal information that facilitates the processing of subsequent foveal information when exploring visual scenes.

### 4.3. Repetition suppression may be achieved by sharpening of the neural code of the object category

There is evidence that object category information of parafoveal images can be extracted within 200 ms after fixation onset ^4^, and in the current study we demonstrated that this information may support to the subsequent processing of foveal images. When an object was primed parafoveally, there was an increase in object category decoding ability around 175 ms after stimulus onset of the foveal image. This indicated that even though the event-related neuronal activity is reduced at this time, the representational specific neuronal information is more specific to the object category. This aligns with our hypothesis that parafoveal priming results in sharpening of the object representation. We however did not find robust support for the theory that gamma oscillations facilitate the inhibitory mechanisms in sharpening ^13,14^. In our study, induced gamma oscillations did not show a significant modulation by parafoveal priming. One explanation for this could be that gamma oscillations might only be sensitive to stimulus repetitions at the same location. In macaque primary visual cortex, gamma power increases occurred only when stimulus repetition was within the receptive fields of the cells ^13^. Unlike previous studies using multiple repetitions of gratings or cropped object images ^13-15^, we repeated the natural images only once. In these studies, the effects on early repetitions on gamma power were varied and did not show a consistent link to sharpening. This, combined with differing stimulus locations, may explain the absence of effects in the gamma band in our parafoveal priming paradigm.

### 4.4. The effects of parafoveal priming are diminished by intervening stimuli

Since we often saccade to similar locations multiple times when scanning a visual scene by eye-movements ^30,31^, our task included a “2-back prime” condition, where intervening images separated the parafoveal prime from the foveal object. This simulated a scenario where parafoveal information primes the foveal object two saccades later, allowing us to assess whether immediate priming effects from the “1-back prime” condition persisted after two simulated saccades. Overall, our results indicate that the presentation of intervening stimuli diminished any parafoveal priming effects, which is in line with results from previous studies ^13,32^. Specifically, in the 2-back prime condition, there was no behavioural benefit on the memory for the foveal items, the repetition suppression effect was greatly reduced, and there was no effect on the object category decoding accuracy. Our “1-back prime” and “2-back prime” condition differed in both the information being presented between the prime and the foveal object and the lag between the prime and foveal object presentation. Therefore, we are unable to comment on what contributed most to the diminishing of the priming effects. It could be the interference from intervening images, the temporal decay in repetition effects, or a combination of both. Additionally, our findings may be linked to inhibition of return (IOR), a mechanism that suppresses attentional priority for previously attended locations ^33^. IOR typically begins around 225 ms and can persist for several seconds ^33,34^.

### 4.5. Conclusion: Parafoveal previewing is associated with sharpening

In conclusion, our study provides evidence that parafoveal priming is associated with both repetition suppression and a more specific neural code, approximately 175 ms after the onset of foveal object processing. The combination of an overall reduction in neural response and the increase in object-specific neuronal code when an object is primed parafoveally supports the idea that parafoveal previewing is associated with sharpening. Our results offer a mechanistic framework for understanding how parafoveal previews may enable efficient visual scene exploration. We argue that parafoveal previewing being associated with sharpening, may support the accumulation of evidence for visual objects when exploring visual scenes.

## 5. Acknowledgements

We would like to thank Sara Calzolari, Camille Fakche, Oscar Ferrante, Brandon Ingram, Dongwei Li, Nina Salman, Yali Pan, Christopher Townsend, Jonathan Winter, Lijuan Wang, and the University of Birmingham’s BlueBEAR HPC service.

This work was also supported by a Wellcome Trust Discovery Award (grant number 227420) and the NIHR Oxford Health Biomedical Research Centre (NIHR203316) attributed to O.J.

## 6. Author Contributions

S.W.: Conceptualization, Methodology, Software, Formal analysis, Investigation, Data Curation, Writing - Original Draft, Writing - Review & Editing, Visualization, Project administration Y.B.: Methodology, Software O.J.: Conceptualization, Resources, Writing - Review & Editing, Supervision, Project administration, Funding acquisition

## 7. Data and Code Availability

Raw data are available upon request, and code can be accessed at [URL will be provided later].

## Notes

### Competing Interest Statement

The authors have declared no competing interest.

